# Motion perception in the common marmoset

**DOI:** 10.1101/522888

**Authors:** Shaun L. Cloherty, Jacob L. Yates, Dina Graf, Gregory C. DeAngelis, Jude F. Mitchell

## Abstract

Visual motion processing is a well-established model system for studying neural population codes in primates. The common marmoset, a small new world primate, offers unparalleled opportunities to probe these population codes in key motion processing areas, such as cortical areas MT and MST, because these areas are accessible for imaging and recording at the cortical surface. However, little is currently known about the perceptual abilities of the marmoset. Here, we introduce a paradigm for studying motion perception in the marmoset and compare their psychophysical performance to human observers. We trained two marmosets to perform a motion estimation task in which they provided an analog report of their perceived direction of motion with an eye movement to a ring that surrounded the motion stimulus. Marmosets and humans exhibited similar trade-offs in speed vs. accuracy: errors were larger and reaction times were longer as the strength of the motion signal was reduced. Reverse correlation on the temporal fluctuations in motion direction revealed that both species exhibited short integration windows, however, marmosets had substantially less non-decision time than humans. Our results provide the first quantification of motion perception in the marmoset and demonstrate several advantages to using analog estimation tasks.

## Introduction

The study of visual motion processing in the primate brain has received considerable attention as a model system for studying neural population codes. This is because the functional neuroanatomy has been well characterized (Maunsell and Vanessen, 1983); (Movshon and Newsome, 1996), stimuli can be easily parameterized, and the motion processing areas of the primate brain have neurons with response properties that are well matched to perceptual features of motion (Born and Bradley, 2005) as well as the sensitivity of psychophysical observers on simple motion discrimination tasks (Britten et al., 1992; Purushothaman and Bradley, 2005; Cohen and Newsome, 2008). As such, motion processing has proven to be a fertile paradigm for studying population codes (Jazayeri and Movshon, 2007b, a; Beck et al., 2008) and even higher cognitive processes such as learning and decision making (Gold and Shadlen, 2007; Law and Gold, 2009).

However, our understanding of the neural code at the circuit level has been hampered by the animal models available for study. One obstacle for circuit level manipulation and measurement in the macaque is that key areas of interest, areas MT and MST, lie buried within the superior temporal sulcus (STS), thus limiting the utility of techniques such as two-photon calcium imaging or large-scale array recordings. At the same time, the neural representation for motion processing in rodents, where those techniques have been well developed, do not appear to involve comparable neural circuits. In contrast to primates, direction selectivity in rodents has a substantial retinal source (Hillier et al., 2017; Shi et al., 2017), areas homologous to MT/MST have not been identified, and psychophysical behavior is remarkably insensitive to large moving stimuli (Marques et al., 2018).

The marmoset monkey offers a potential opportunity for studying neural population codes that underlie motion processing using modern methods. Recently, the marmoset monkey has emerged as a model for visual systems neuroscience (Mitchell et al., 2014; Mitchell et al., 2015; Johnston et al., 2018) that may overcome several long-standing limitations of other primate species. The marmoset has established homologies with the macaque and the human brain (Solomon and Rosa, 2014), including quantitative similarities in motion processing circuitry and function (Lui and Rosa, 2015). Importantly, unlike in macaques or humans, most cortical areas in the marmoset brain lie on the surface and are readily accessible for recording (Solomon et al., 2011; Sadakane et al., 2015; Solomon et al., 2015; Zavitz et al., 2016; Zavitz et al., 2017). At present, there is a concerted effort by several international groups to develop transgenic marmoset models (Sasaki et al., 2009; Izpisua Belmonte et al., 2015; Okano et al., 2016), providing novel molecular tools, such as gCaMP6 lines for use in two photon imaging (Park et al., 2016), as well as genetic models of human mental disease (Okano et al., 2016).

While the neuroanatomy and basic sensory processing of visual motion stimuli have been studied in marmosets (Solomon et al., 2011; Zavitz et al., 2016; Chaplin et al., 2017), little is known about their perceptual abilities. Here, we introduce a novel motion estimation task that is ideally suited for studying motion perception in marmosets. Two marmosets were trained to indicate their perceived direction of motion by making a saccadic eye movement to a “target ring” that surrounded the motion stimulus. Beyond the utility for training, estimation tasks can offer more information about the perceptual process than traditional classification or discrimination paradigms (Pilly and Seitz, 2009) and have been useful for studying readout mechanisms for motion perception (Nichols and Newsome, 2002; Jazayeri and Movshon, 2007b, a; Webb et al., 2007; Webb et al., 2011). The trial-by-trial distribution of responses from an estimation task supports a much richer description of the underlying perceptual process than binary reports from traditional 2AFC paradigms (Laquitaine and Gardner, 2018).

We show that the precision of the marmosets’ perceptual reports varies systematically with the strength of the motion signal in a similar manner to human observers. We then compare the performance of the marmosets to that of human observers performing the same estimation task and, using behavioral reverse correlation, directly quantify both marmoset and human temporal integration properties. In general, marmoset behavior closely resembled human performance, except that marmosets were less precise, had a larger dependence of reaction time on task difficulty, and required less time to plan an eye movement than human subjects. Taken together with known physiological properties of motion processing areas in marmosets, these results establish the marmoset monkey as a viable model system for human motion perception and for studying the underlying neural population codes.

## Material and Methods

Data were collected from two adult male marmoset monkeys (*Calithrix jacchus*) and four human psychophysical observers. All surgical and experimental procedures involving the marmosets were approved by the Institutional Animal Care and Use Committee at the University of Rochester, and by the Animal Ethics Committee at Monash University. All procedures involving human psychophysical observers were approved by the Research Subjects Review Board at the University of Rochester.

### Surgical procedures

Marmosets were implanted with a titanium head-post to stabilize their head during behavioral training. Surgical procedures were performed under aseptic conditions and were identical to those described previously (Nummela et al., 2017).

### Visual stimuli and behavioral training

Visual stimuli were generated in Matlab (The Mathworks, Inc.) and presented using the Psychophysics toolbox (Brainard, 1997) at a frame rate of 120 Hz on a LCD monitor (XT2411z; BenQ) placed 60 cm in front of the animals. The monitor had a mean luminance of 115 cd/m^2^ and resolution of 1920 × 1080 pixels (W × H) covering 48° × 28° (W × H) of visual angle.

Marmosets sat in a purpose-built chair (Remington et al., 2012) with their head fixed by way of the implanted head-post. After habituating to being head-fixed, both marmosets were trained to maintain fixation within a small window around a fixation target presented at the center of the screen. Marmosets received liquid reward for maintaining fixation on this central target. After fixation training, but prior to training on the motion estimation task, both animals were trained to make visually guided center-out saccades in a grating detection task (Nummela et al., 2017; monkey S also participated in our previous study, see Subject S).

Both subjects were then initially trained to perform a coarsely discretized version of the motion estimation task (Fig. 1A). To initiate each trial, the marmoset was required to maintain fixation within a small window (radius 1.8°) around a fixation target presented in the center of a uniform gray screen of mean luminance. This fixation target consisted of two concentric black (0.5 cd/m^2^) and white (230 cd/m^2^) circles, 0.3° and 0.6° in diameter, respectively. After a prescribed fixation period of 200–500 ms (drawn randomly on each trial, from a uniform distribution), a random pattern of dots (40 dots each 0.2° in diameter; luminance 97 cd/m^2^) was presented within a circular aperture 7° in diameter, centered on the fixation target. During initial training, the dots moved coherently at a speed of 15 °/s in one of 8 possible directions, drawn randomly on each trial from the set {0°, 45°, …, 315°}. Dots had a limited lifetime of 50 ms (6 frames). At the start of each trial, each dot was assigned an age (in frames) drawn randomly from a uniform distribution between 0 and 6 frames. At the end of their lifetime, dots were redrawn at a random position within the stimulus aperture and their age was reset to zero. Dots that exited the aperture were replaced on the opposite side of the aperture.

**Figure 1.**
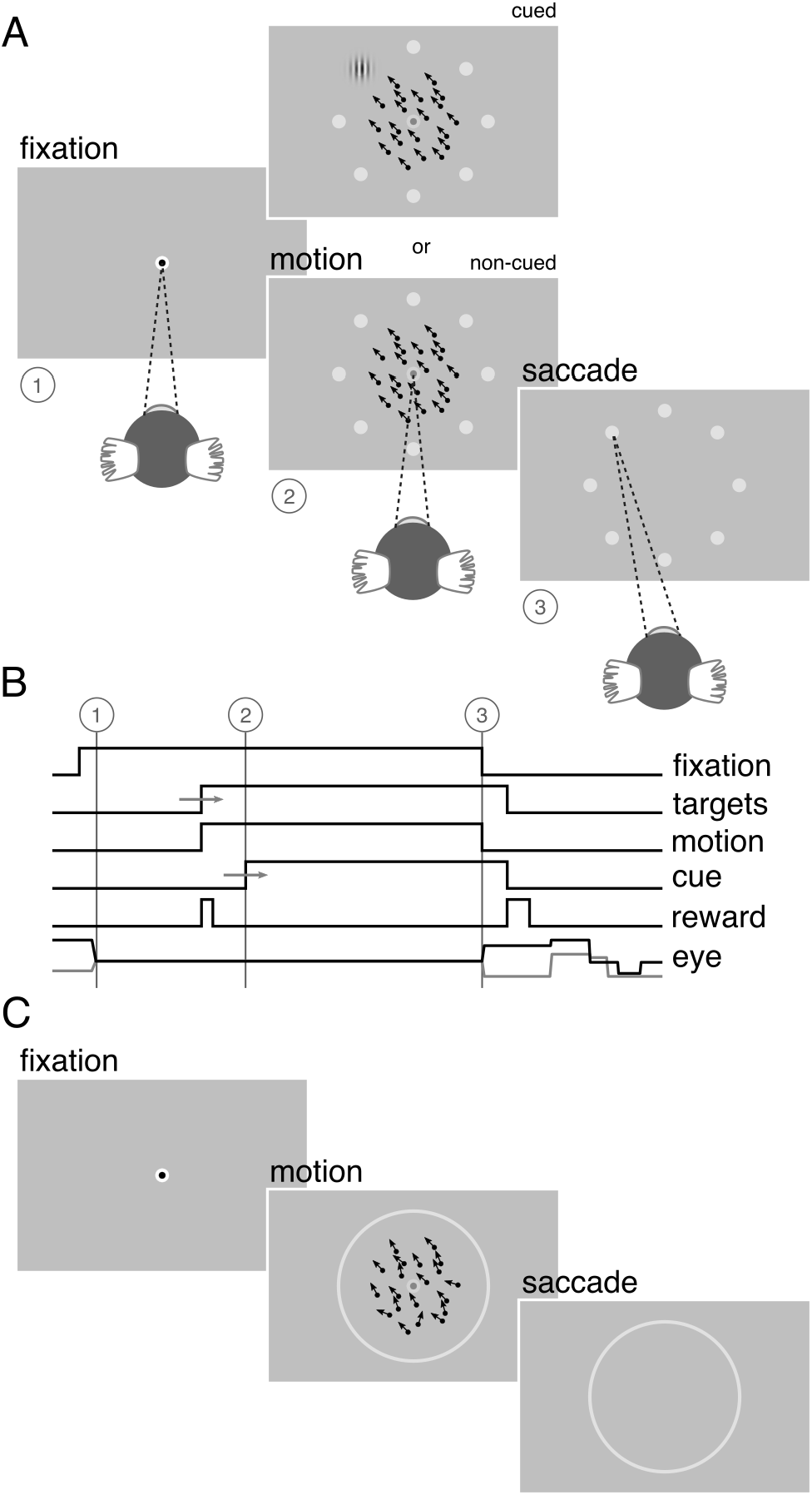
Visual stimuli and behavioral task. **A**. Marmosets were trained to maintain fixation within a window 2° in diameter around a target presented at the center of the screen (1). A random pattern of dots was then presented within a circular aperture 7° in diameter centered on the fixation target (2). The dots moved at a speed of 15°/s in one of 8 possible directions equally distributed between 0–360°. Coincident with the onset of the random dot pattern, 8 small choice targets were presented, equally spaced around a ring, 10.6° in diameter, concentric with the central fixation target. Marmosets received a liquid reward for correctly reporting the direction of motion by making a saccade to one of the choice targets (3). On a proportion of trials, marmosets received an overt cue, consisting of a small high contrast Gabor patch, presented at the location of the correct choice target. On these trials, marmosets could obtain the reward by making an eye movement to the cued target location without integrating the motion stimulus. These cued trials served to ensure a sufficiently high rate of reward to keep the marmosets engaged with the task. Over the course of training, the proportion of cued trials was gradually reduced. **B**. Sequence of trial events. After a fixation period of 200–500 ms (1), the random dot pattern appeared (2). Marmosets were required to maintain fixation on the central target for a minimum duration of 100 ms after appearance of the motion stimulus, after which the fixation target dimmed and the marmosets were free to indicate the perceived direction of motion by making an eye movement to one of the choice targets. Both the fixation point and the random dot pattern were extinguished if the marmoset broke fixation or after a maximum period of 600 ms, whichever came first (3). **C**. After initial training, the number of possible motion directions was increased from 8 to 50 over the course of several weeks and the discrete choice targets were replaced by a continuous ring. The strength of the motion signal was then varied by assigning to each dot a direction drawn from a uniform generating distribution centered on the target motion direction.

The marmoset was required to maintain fixation on the central target for a minimum duration of 100 ms after onset of the motion stimulus, after which the fixation target dimmed, signaling that the marmoset was free to indicate the perceived direction of the motion by making an eye movement to one of 8 possible choice targets. The choice targets were small gray (120 cd/m^2^) circles, 0.6° in diameter, presented at 8 equally spaced points, {0°, 45°, …, 315°}, around a circle 10.6° in diameter centered on the central fixation target. One choice target corresponded to each of the possible motion directions. Task timing is illustrated schematically in Figure 1B. To discourage guessing as a strategy, we imposed a minimum interval between trials of ≥2 s, during which the monitor displayed a uniform gray screen of mean luminance.

Marmosets received a small liquid reward (typically 5–10 μl) for initiating a trial (i.e., after the initial fixation period, before onset of the motion stimulus) and again at the end of the trial for correctly reporting the direction of motion. The volume of the latter reward was scaled according to the angular error in the marmoset’s choice such that larger errors received smaller rewards. Errors less than 8° from the true motion direction received the maximal reward (typically 20–40 μl) while errors greater than 27° were unrewarded (i.e., 0 μl). This reward schedule served to ensure a sufficient rate of reward to keep the marmosets engaged with the task. Marmosets also received visual feedback presented at the location of the correct choice target. Correct choices were indicated by presentation of a small marmoset face. Incorrect choices were indicated by presentation of a dark gray (57 cd/m^2^) circle 2.7° in diameter surrounding the correct choice target.

During the training period, on a proportion of trials, marmosets received an overt cue as to the correct choice target. This cue consisted of a small high contrast Gabor patch (100% contrast, 4 cycles/°, 2.7° in diameter) presented at the location of the correct choice target. On these cued trials, marmosets could obtain the reward by making an eye movement to this cue, without regard to the presented motion direction (although the motion direction was always predictive of the cue and the reward). During initial training, the cued trials served to ensure a sufficiently high rate of reward to keep the marmosets engaged with the task and help establish the association between the motion direction, the choice targets, and the reward. Over the course of training, the proportion of cued trials was gradually reduced and eventually eliminated by delaying the time window during which the cue appeared, encouraging the marmosets to anticipate its location based on the direction of motion presented.

### Behavioral task

After the initial training regimen described above, the difficulty of the estimation task was manipulated in two ways (Fig. 1C). First, the number of possible motion directions was progressively increased, over several weeks, until the motion direction was selected from 50 possible directions, drawn randomly on each trial, from the set {0°, 7.2°, …, 352.8°}. Once the number of possible motion directions exceeded 18, the discrete choice targets were replaced with a continuous ring, 10.6° in diameter, drawn with the same luminance as the original choice targets. In this configuration, the marmosets’ behavioral reports were continuous around this ring, similar to the motion estimation task described by Nichols and Newsome (2002). Second, we then varied the difficulty of the task by adjusting the range of dot directions on each trial by sampling the direction assigned to each dot from a uniform generating distribution. The mean of this generating distribution defined the target motion direction (drawn randomly on each trial from the set of 50 possible motion directions) and the width of the generating distribution defined the range of individual dot motion directions. This manipulation is similar to that described by (Zaksas and Pasternak, 2006). The width, or range, of the generating distribution was drawn randomly on each trial from the set {0°, 45°, 90°, …, 360°}. In the analyses below, we describe behavioral performance as a function of signal strength, *SS*, given by *SS* = 1 − Range/360. *SS* = 1 corresponds to coherent motion while *SS* = 0 corresponds to random, incoherent motion. To ensure a sufficient rate of reward and keep the marmosets engaged with the task, the more difficult conditions, *SS* ≤ 0.5, were presented on only 35% of trials while the remaining 65% of trials were assigned *SS* > 0.5.

### Recording and analysis of eye position

Eye position was sampled continuously at 220 Hz using an infrared eye tracker (USB-220, Arrington Research). Methods for calibrating the eye tracker in each daily session were identical to those described previously (Nummela et al., 2017). This calibration procedure set the offset and gain (horizontal and vertical) of the eye tracking system. To mitigate any uncalibrated rotational misalignment (around the optical axis) of the eye tracking camera, which would be manifest in our data as a non-zero bias in subjects’ errors, we computed the mean angular error over all trials of the motion estimation task for each subject and subtracted this rotational component prior to further analysis.

All analyses were performed off-line in Matlab (The Mathworks, Inc.). Saccadic eye movements were identified automatically using a combination of velocity and acceleration thresholds. First, the raw eye position signals were resampled at 1 kHz, and horizontal and vertical eye velocity signals were calculated using a finite impulse response digital differentiating filter (Matlab function lpfirdd() (Chen, 2003) with parameter N = 16 and a low-pass transition band of 50–80 Hz; this filter has a −3 dB passband of 19–69 Hz). Horizontal and vertical eye acceleration signals were calculated by differentiation of the velocity signals using the same differentiating filter. Negative going zero crossings in the eye acceleration signal were identified and marked as candidate saccades. These points correspond to local maxima in the eye velocity signal. Eye velocity and acceleration signals were then examined within a 150 ms window around each candidate saccade. Candidate saccades were retained provided that eye velocity exceeded 10 °/s and eye acceleration exceeded 5000 °/s^2^. Saccade start and end points were determined as the point preceding and following the peak in the eye velocity signal at which eye velocity crossed the 10 °/s threshold.

Drift in eye position during presentation of the motion stimulus was quantified as follows. Raw horizontal and vertical eye position signals were first smoothed with a median filter (Matlab function medfilt1() with parameter N = 3; at the sampling frequency of 220 Hz this filter has a low-pass characteristic with a −3 dB cut-off frequency of ~50 Hz) to minimise high frequency noise from the eye tracking camera. The smoothed eye position signals were then resampled at 1 kHz, and horizontal and vertical eye velocity signals calculated using a finite impulse response digital differentiating filter (lpfirdd() (Chen, 2003) with parameter N = 16 and a low-pass transition band of 30–50 Hz; this filter has a −3 dB passband of 9–49 Hz). Segments beginning 20 ms before and extending until 20 ms after each saccadic eye movement (identified as described above) were then removed, to minimize saccadic intrusion in the drift velocity estimates. Systematic drift in the direction of the motion stimulus was quantified by projecting the horizontal and vertical eye velocity signals onto the stimulus motion direction, adding them, and then averaging the resultant across all trials for each of the motion signal strengths. Missing values, due to removal of saccades, did not contribute to the average across trials. The average eye velocity traces were characterized by a period immediately after motion onset during which the eye remained stationary, followed by a period of increasing drift velocity terminated by the saccade indicating the animal’s choice (Fig. 8B). For both marmosets, the period of drift began approximately 75 ms after onset of the motion stimulus. The magnitude and direction of drift was computed as the vector sum of the horizontal and vertical eye velocity signals beginning 75 ms after onset of the motion stimulus.

### Analysis of behavior

To quantify task performance during training, we computed the angular error between the target motion direction on each trial and the marmoset’s behavioral choice. Behavioral choice was calculated as the angle of the median of the eye position measured in the first 25 ms after entering the acceptance window on the target ring. For the majority of trials, the marmoset’s choices are correlated with the direction of motion presented, and their errors reflect a combination of noise in sensory processing and the motor output. However, to account for trials on which the marmoset’s choices reflect random guesses or momentary lapses in attention, we modeled their errors as random variates drawn from an additive mixture of two probability distributions: a wrapped Normal distribution (reflecting errors in perceptual processing of the motion stimulus or in motor execution) and a uniform distribution (reflecting non-perceptual errors or ‘lapses’). This mixture model is described by three parameters: λ, the height of the uniform distribution, representing the lapse rate, and the mean, μ, and standard deviation, σ, of the wrapped Normal distribution. The mean (μ) represents any systematic bias in the marmoset’s choices away from the target motion direction. Because performance on this task is described by deviations from the true direction, we fixed μ to zero for our analyses, and we quantify performance by the standard deviation (σ) of the wrapped Normal distribution. The parameter σ represents the performance of the subjects after accounting for lapses, and captures both perceptual and motor noise. We identified the parameters λ and σ for each session by maximizing the likelihood of the observed errors.

To investigate the ability of marmosets to estimate the direction of noisy motion, we examined the distribution of errors in perceptual choices over a range of motion signal strengths, as described above. For this purpose, we again fit the mixture model just described, pooling trials across all sessions.

On each trial we also recorded the marmosets’ reaction time as the interval from onset of the motion stimulus until the marmosets reported their choice. In all analyses we estimated either standard errors (s.e.) or 95% confidence intervals, via a bootstrap sampling procedure (Efron and Tibshirani, 1993).

### Psychophysical reverse correlation

To estimate each subject’s temporal weighting function, we correlated the errors in their behavioral reports with the random fluctuations in stimulus direction that arise from the underlying generating distribution and the limited lifetime of the dots. Specifically, we computed temporal weighting functions, or kernels, in two ways: first, with all trials aligned at the time of motion stimulus onset, and second, after realigning all trials at the onset time of the saccade indicating the subject’s choice. In each case, we computed Pearson’s linear correlation, R, between the mean dot direction on each stimulus frame and the subject’s perceptual choices, across all trials. The resulting kernels reveal the contribution of each stimulus frame to the subject’s perceptual choices.

From the saccade-aligned kernels, we estimated each subject’s dead time – the time required to plan an eye movement after reaching their decision – by fitting a single-knot linear spline to the saccade-aligned kernel on the interval [*t*_pk_, 0], where *t*_pk_ denotes the time preceding saccade onset corresponding to the peak of the temporal kernel. Specifically, we defined the piecewise linear spline function

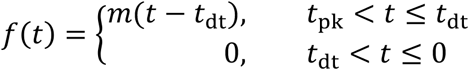

and fitted the slope, *m*, and the dead time (*t*_dt_; constrained such that *t*_pk_ < *t*_dt_ ≤ 0) by minimizing the sum of squared residuals between the spline, *f*(*t*), and the saccade aligned kernel amplitude.

### Human psychophysics

To compare the marmosets’ performance with that of human observers, we had four human subjects (two female and two male, ages 21-46 years; including two of the authors JLY and JFM (Humans 2 and 4, respectively), and two naïve subjects) perform the same motion estimation task. Each subject performed at least four (4) sessions, on separate days. Two subjects (Humans 1 and 2) underwent more extensive training (10 and 8 sessions, respectively). Subjects had normal or corrected to normal vision and sat comfortably with their head immobilized by way of a bite bar. All other equipment for experiment control, stimulus presentation, and eye tracking was identical to that used for the marmoset experiments. All task and stimulus parameters for the human subjects were also identical to those described above for the marmosets, with the exception that human subjects received instruction on the requirements of the task, received auditory feedback rather than liquid reward, and did not receive an overt cue on any trials. Auditory feedback consisted of 1–4 clicks based on the angular error: errors < 8° produced 4 clicks and errors >27° produced no auditory feedback.

## Results

### Initial task training

We trained two marmosets to perform the motion estimation task (monkey S, 68 sessions over 166 days; monkey H, 44 sessions over 92 days; Fig. 2). Initially, marmosets initiated relatively few trials (~100 trials/session for both monkey S and monkey H; Fig. 2A, open symbols). However, within 1–2 sessions both marmosets learned the requirements of the task and, over the course of training, reliably initiated several hundred trials per session on average (mean ± s.e.m, 406 ± 17.9 trials for monkey S; 325 ± 20.7 trials for monkey H). Of the trials initiated, both marmosets initially completed only a small fraction (Fig. 2A, filled symbols). A trial was deemed to be complete if the marmoset maintained fixation for at least 100 ms after onset of the random-dot motion stimulus (see Fig. 1), made a saccade of >3° out of the fixation window and towards the choice targets, and maintained that new fixation location for at least 25 ms. For both marmosets, the number of completed trials increased very quickly initially (e.g., within 2–3 sessions), and then more slowly thereafter. Over the course of training, both marmosets completed approximately 100 trials per session, on average (mean ± s.e.m, 123 ± 6.0 trials/session for monkey S; 108 ± 11.1 trials/session for monkey H). As a proportion of the total trials initiated, monkey S showed a small increase in trial completion rate over the course of training, while for monkey H completion rate remained approximately constant. Much later in training (> 1 year), we withheld the reward delivered after initial fixation and reduced the inter-trial interval from ≥2 s to ≥0.5 s. On average both marmosets then completed a greater number of trials within each session (Fig. 2A). This increase was most evident for monkey H (mean ± s.e.m, 200 ± 32.2 trials/session for monkey S; 340 ± 20.6 trials/session for monkey H).

**Figure 2.**
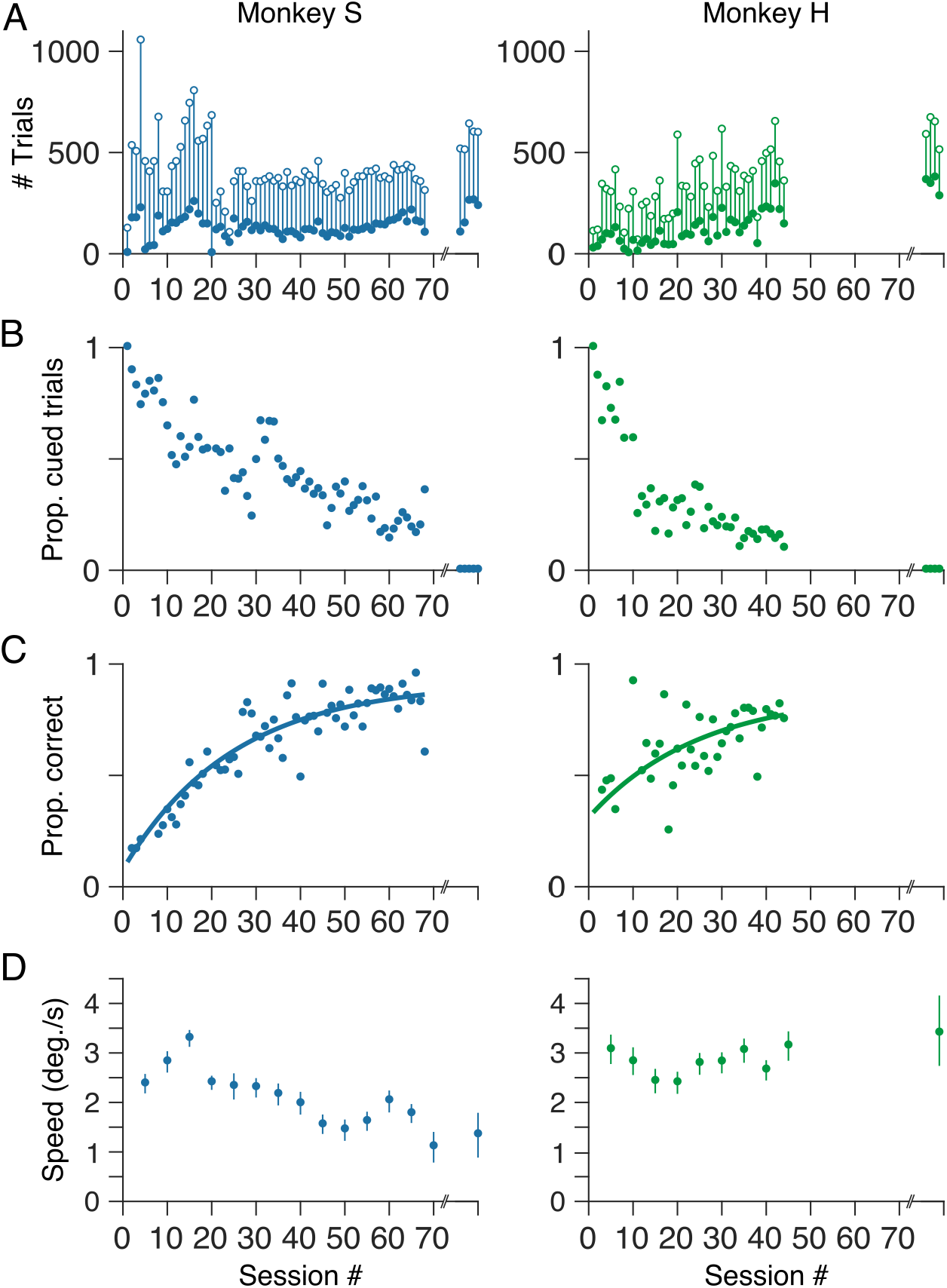
Initial task training. **A.** Total number of trials (open symbols) together with the number of completed trials (filled symbols) per session. Trial counts over one week (5 sessions for monkey S; 4 sessions for monkey H) after approximately 1 year of training are shown on the right in each panel. In these sessions, the monkeys were performing the continuous version of the estimation task with 50 possible motion directions and 9 possible motion signal strengths. **B.** Initially, both marmosets based their choices on the overt cue rather than motion of the random dot pattern. However, over the course of training the proportion of cued trials decreased as the onset of the cue was progressively delayed. **C**. As a measure of performance we computed the proportion correct – the proportion of non-cued trials in each session in which the marmoset’s choice fell within the reward window (see Material and methods). Proportion correct for both marmosets improved over the course of training. Solid lines show least-squares fits of a single exponential function. **D**. Speed of drift in eye position during presentation of the motion stimulus, over the course of training.

As described above, on a proportion of trials early in training, the marmosets received an overt cue as to the direction of motion (i.e., the correct choice target). The time at which this cue appeared, after onset of the motion stimulus, was randomized from trial to trial. We designated each complete trial as either *cued* or *non-cued*, depending on whether the cue appeared before or after the marmoset indicated their choice. We then computed the number of cued trials, as a proportion of completed trials, within each session over the course of training. Initially, both marmosets relied heavily on the overt cue (Fig. 2B). However, with training both marmosets learned the association between the direction of motion of the random-dot pattern and the correct choice target, and to base their choices on their perception of motion direction rather than wait for the overt cue (Fig. 2B). We encouraged this behavior, over the course of training, by progressively delaying the temporal window within which the cue appeared until the cue was not presented at all.

### Motion estimation improves with training

To quantify task performance during training, we considered only the non-cued trials within each session and computed the proportion of trials (within each session) in which the monkey’s choices fell within the reward window (i.e., within ±28° of the target motion direction; see *Materials and methods*). For both marmosets, the proportion of correct (i.e., rewarded) choices increased over the course of training (Fig. 2C). Proportion correct as a function of session was fit by a single exponential function with an upper asymptote of 0.92 ± 0.02 for monkey S and 0.87 ± 0.06 for monkey H (time constant: 24.8 ± 1.6 sessions for monkey S, 24.9 ± 5.7 sessions for monkey H).

### Stimulus-dependent eye drift from fixation is reduced with training

Several aspects of our paradigm were designed to minimize pursuit eye movements evoked by the motion stimulus, and to discourage the animals from pursuing the stimulus for reward. Specifically, the central fixation point remained on the screen (although at reduced contrast) for the duration of the motion stimulus, the motion stimulus itself was of moderate contrast, and the dots had limited lifetime (see *Materials and methods*). Nevertheless, we often observed small drifts in eye position during fixation, evoked by presentation of the motion stimulus (for an example, see Fig. 8A). These movements were brief and of low gain relative to the stimulus velocity, typically less than 20–30% gain (less than 3 degrees of visual angle per second). During the brief periods prior to the saccade choice this drift resulted in displacements in eye position that were typically less than 0.5 degrees of visual angle, which was not sufficient to break the task defined fixation window. To quantify the magnitude of the stimulus evoked drift we projected instantaneous eye velocity (see *Materials and methods*) onto the direction of the motion stimulus, and averaged the resulting eye speed signals across trials. To quantify any systematic variation over the course of training we pooled trials across five sessions at a time. We then computed the average eye speed over a 50 ms window beginning 200 ms after onset of the motion stimulus. This window corresponded to the peak in the average eye speed signal for both marmosets (for example eye speed signals, see Fig. 8B). Early in training, both marmosets showed average drift speeds of approximately 3 deg./s (Fig. 2D). For monkey S, drift speed decreased over the course of training (Fig. 2D; left), likely reflecting improved fixation control. Average drift speed for the second marmoset (monkey H) remained relatively constant throughout training. (Fig. 2D; right). Notably, both marmosets exhibit small drifts in eye position even after training. Human observers also exhibited similar drifts in eye position, though smaller in magnitude (mean ± s.d., 0.6 ± 0.36 deg./s). After considering the accuracy of saccade choices, we also quantify the accuracy of drift eye movements as a measure of the stimulus motion direction, and compare the precision of the drift to that of the saccade choices.

### After training, choices reflect motion direction

The improvement in motion estimation performance with training is reflected in the distributions of saccade end-points (i.e., choices) for the two marmosets (Fig. 3A). For both marmosets, the distribution of saccade end-points reflects the distribution of directions presented. Specifically, saccade end-points for both monkeys are distributed around an annulus, and are grouped according to target motion direction (like-colored points are grouped, with an orderly progression of colors around the annulus).

**Figure 3.**
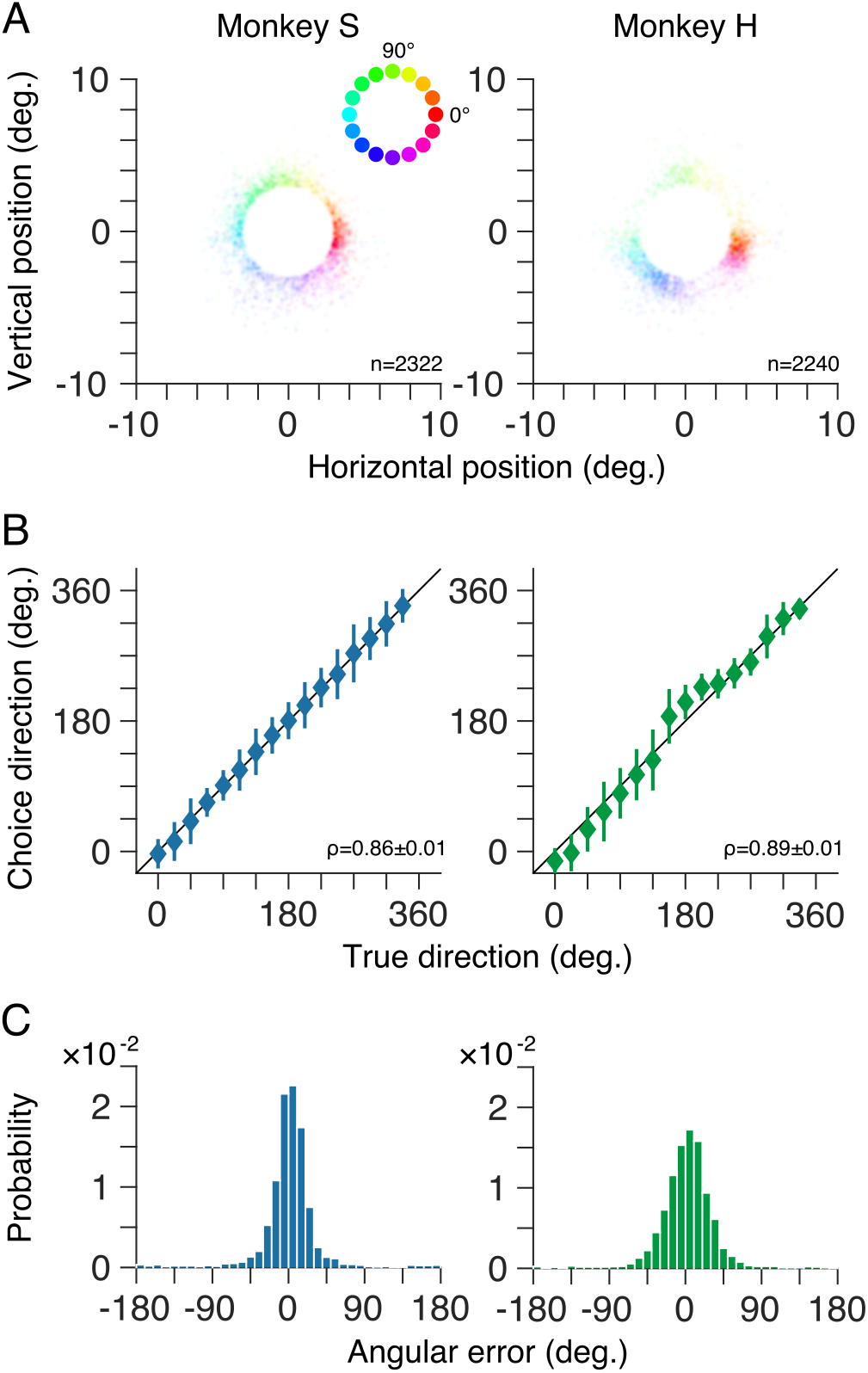
Choices reflect stimulus motion direction after training. **A**. Distributions of saccade end-points for non-cued trials for both marmosets as kernel density plots. At each spatial location, target motion direction is represented by the hue (see inset) while the density of saccade end-points (i.e., trials) is represented by the saturation. For the trials shown, random dot patterns moved coherently in one of 50 possible directions between 0–360°. **B**. Mean choice direction (symbols) for non-cued trials plotted against target motion direction. Behavioral choices of both marmosets were highly correlated with the target motion direction. Error bars show ± 1 standard deviation. **C**. Distributions of angular error, the difference between the marmoset’s choice and the true motion direction on each trial, for both marmosets.

We quantified the performance of both marmosets, after training, in two ways. First, we computed the circular correlation between their choices and the presented target motion direction. The choices of both marmosets were strongly correlated with the target motion direction (Fig. 3B; ρ = 0.86 ± 0.01 for monkey S; ρ = 0.89 ± 0.01 for monkey H; mean ± s.e.m). Second, we computed the distribution of angular errors between the monkey’s choice and the target motion direction on each trial. For both marmosets the distribution of errors deviate significantly from uniform (p < 0.001; Rayleigh test) with unimodal peaks close to 0 (Fig. 3C). The standard deviation of the error distributions, a measure of the precision of the perceptual reports, was 34.4 ± 1.2° for monkey S and 33.1 ± 1.0° for monkey H (mean ± s.e.m).

### Performance increases through learning

The distributions of angular errors are well described by a mixture of a uniform distribution (reflecting non-perceptual errors or ‘lapses’) and a wrapped Normal distribution (reflecting errors in perceptual processing or motor output; see *Material and Methods*; Fig. 4A). We further quantified the marmosets’ performance during training by fitting the mixture model to the distribution of angular errors within each session. The performance of one marmoset (monkey S) was initially very poor but improved over the course of training (Fig. 4B; left). The reduction in standard deviation of the error distribution with training was fit by a single exponential function (solid curve in Fig. 4B; left) with an asymptote of 15.8 ± 1.4° (mean ± s.e.m). The performance of the second marmoset (monkey H) was good even in very early sessions. Estimates of the standard deviation of the error distribution for this marmoset were initially variable (from session to session) but converged, with training, to a value of 25.8 ± 4.7°.

**Figure 4.**
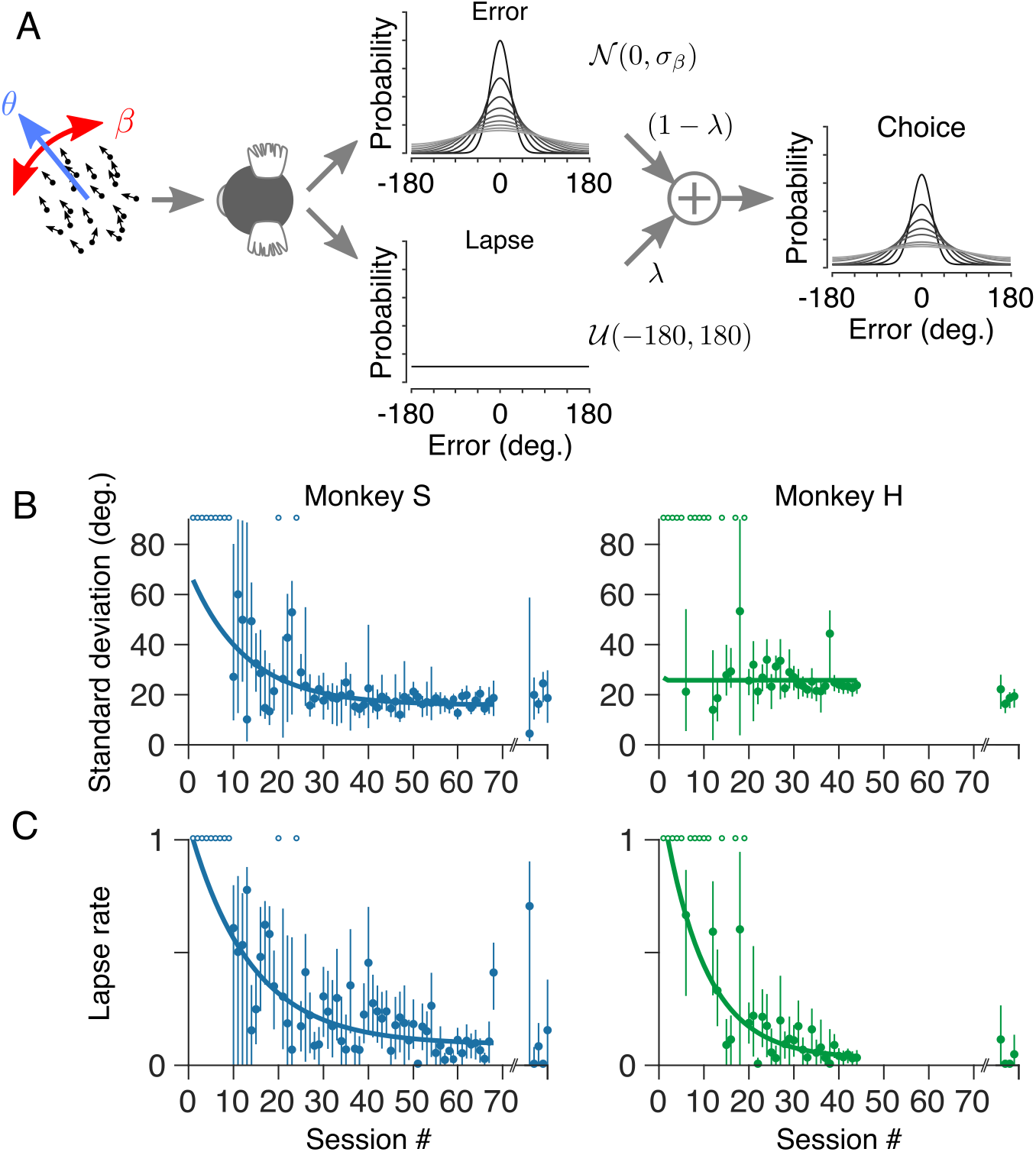
Motion estimation performance improved with training. **A.** To quantify behavioral performance, both during training and subsequently on the main task, we modelled the distributions of angular errors as a mixture of two probability distributions: a uniform distribution (reflecting non-perceptual errors or ‘lapses’) and a wrapped Normal distribution (reflecting errors in perceptual processing of the motion stimulus). The relative contribution of these two distributions is determined by the lapse rate, λ. Task performance is quantified by the standard deviation, σ, of the wrapped Normal distribution. **B**. Standard deviation of both marmosets’ errors plotted as a function of training session number. **C.** Lapse rate plotted as a function of training session number for both marmosets. The lapse rate of both marmosets decreased with training. Note that we were unable to fit the mixture model in some sessions, particularly early in training when the marmosets performed relatively few non-cued trials in each session. Sessions containing too few (< 30) non-cued trials are indicated by open symbols at the top of the axes in B and C. Symbols on the far right of each panel in B and C show standard deviation and lapse rate, respectively, after more than 1 year of training (see Figure 2). Solid curves show least-squares fits of a single exponential function.

The lapse rate of both marmosets decreased with training (Fig. 4C) and was fit by a single exponential function (solid curves in Fig. 4C) with asymptotes of 0.09 ± 0.04 (monkey S; mean ± s.e.m) and 0.03 ± 0.02 (monkey H). Both marmosets therefore learned the task during training but with distinct learning trajectories.

Note that in some sessions, particularly early in training, we could not reasonably fit the mixture model to estimate the standard deviation of the errors or the subject’s lapse rate, either because the available data were insufficient (<30 non-cued trials within a session) or the distribution of errors was inconsistent with the mixture model (e.g., errors better described by a uniform distribution). These sessions are indicated by open symbols at the top of each panel in Figure 4B and 4C.

### Accuracy and reaction time vary systematically with signal strength

During training (sessions 1–68 for monkey S, and sessions 1–44 for monkey H; Figs. 2–4), marmosets saw only stimuli with coherent motion (i.e., signal strength, *SS* = 1.0). After the monkeys learned the task, as evidenced by significant correlations between choice and motion direction (Fig. 3), and by a plateau in their performance (Fig. 4), we randomly interleaved trials with a range of motion signal strengths. We varied signal strength by varying the width of the generating distribution. The smallest width of the generating distribution (0°) assigned all dots the same motion direction (*SS* = 1). Greater widths of the generating distribution introduced greater direction scatter among dots, with the largest possible width (360°) reflecting uniform sampling of all possible motion directions (*SS* = 0). After training, marmosets performed this motion estimation task in daily sessions over several months (17,021 trials over 85 sessions for monkey S, and 17,937 trials over 77 sessions for monkey H). On average, the marmosets performed the task for 20–30 minutes per day (mean ± s.d., 25.7 ± 4 minutes for monkey S; 26.7 ± 4 minutes for monkey H). To investigate how marmosets pool motion signals to make perceptual decisions, we computed psychometric and chronometric functions for both marmosets. These data consisted of the angular error of the marmoset’s choice (Fig. 5; see *Materials and methods*) and their reaction time (the interval from onset of the motion stimulus until the marmoset indicated a choice; Fig. 6) for each completed trial, for all target motion directions and all motion signal strengths.

**Figure 5.**
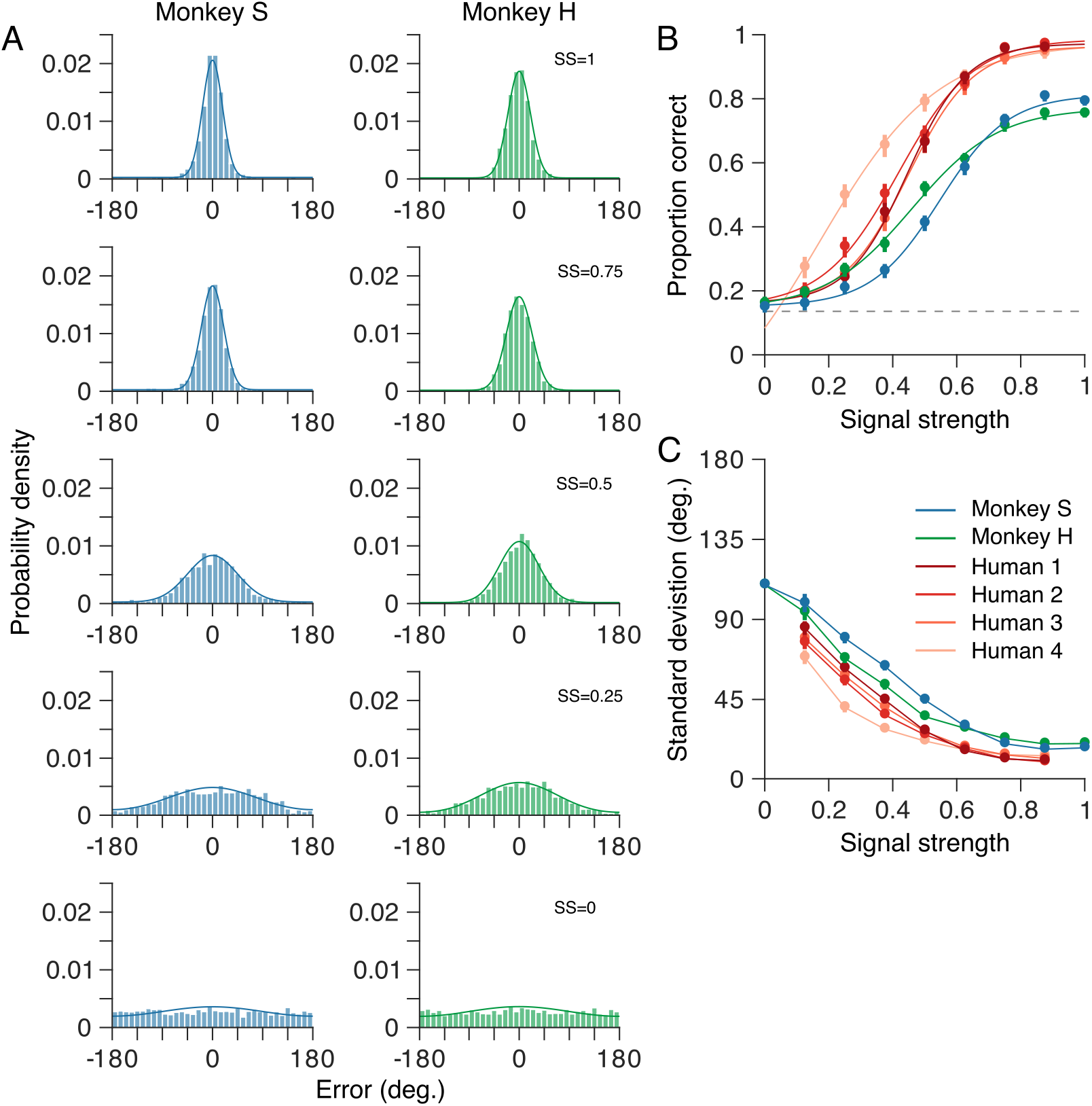
Perceptual errors vary systematically with motion signal strength. **A**. Distributions of angular errors – the difference between the marmoset’s choice and the true motion direction on each trial – for a range of motion signal strengths (see Material and methods). Signal strength, SS = 1, corresponds to coherent motion while SS = 0 corresponds to random, incoherent motion. Error distributions (bars) of both monkeys became broader as signal strength was reduced. The data include 17,021 trials, across all conditions, from 85 sessions for monkey S and 17,937 trials from 77 sessions for monkey H. Solid curves show the probability density defined by the mixture model (see Materials and methods) fitted to the error distributions. **B**. Proportion of correct (i.e., rewarded) trials as a function of signal strength. Proportion correct for both marmosets decreased as signal strength was reduced. In the absence of any coherent motion signal (SS = 0), both marmosets performed at the chance level (dashed line). **C**. Standard deviation of the mixture model, fitted to each marmoset’s errors, as a function of stimulus strength. The standard deviation of both marmosets’ errors increased as signal strength was reduced. For comparison, B and C show comparable metrics for four human observers performing the same motion estimation task (see Material and methods). The human observers performed better than the marmosets over all signal strengths. In B and C, error bars show bootstrap estimates of the 95% confidence interval for the corresponding metric. Solid curves in B show maximum likelihood fits of a logistic function.

**Figure 6.**
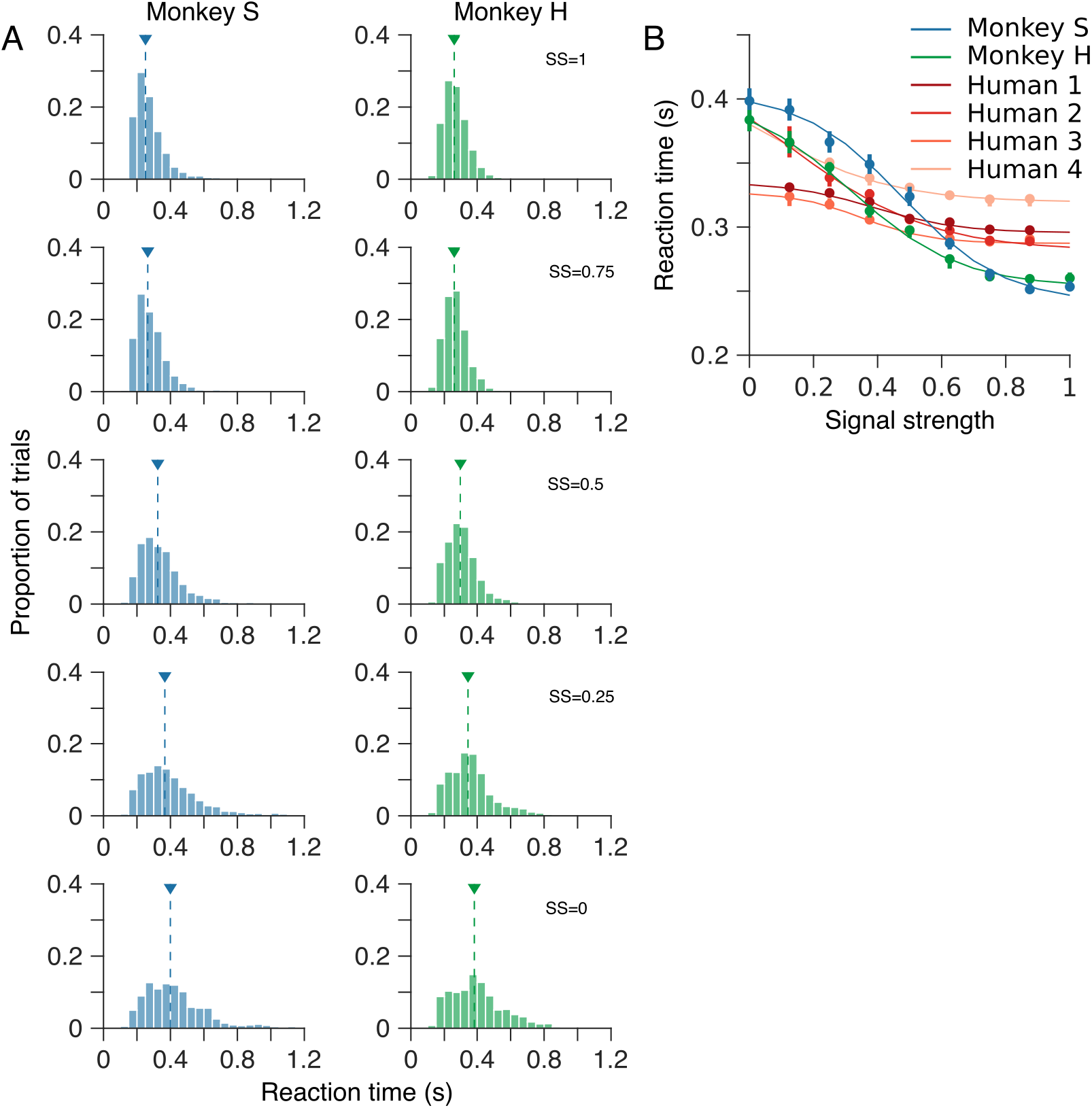
Reaction time varies systematically with motion signal strength. **A**. Distributions of reaction time – the time interval from onset of the motion stimulus until the marmosets’ indicated their choice – for a range of motion signal strengths. Arrow heads indicate the median reaction time for each distribution. **B**. Median reaction time as a function of signal strength. For comparison, B also shows median reaction times for four human observers performing the same motion estimation task. Both humans and marmosets exhibit a typical increase in reaction time as signal strength was reduced. However, the increase in reaction time of the human observers was less dramatic than that of the marmosets. In B, error bars show bootstrap estimates of 95% confidence intervals. Solid curves show least-squares fits of a hyperbolic tangent function.

Figure 5A shows distributions of angular errors for both marmosets over a range of motion signal strengths. We quantified their performance in two ways. First, we computed the proportion of trials on which their choices fell within the reward window (i.e., error < 28°; see *Materials and methods*). Second, we fitted the mixture model to the distributions of angular errors, pooling the data across all conditions (i.e., all target motion directions and all motion signal strengths). Specifically, we fixed the lapse rate, λ, across all signal strengths but allowed the standard deviation of the wrapped Normal distribution to vary with signal strength.

For strong motion signals, the proportion correct for both marmosets approached or exceeded 0.8 (proportion correct 0.83 and 0.79, 95% CIs [0.82, 0.84] and [0.77, 0.80], for monkeys S and H, respectively; Fig. 5B), and the standard deviation of their errors, as quantified by the mixture model, was similar (17.9° and 20.3°, 95% CIs [17.2, 18.5] and [19.8, 21.0], for monkeys S and H, respectively; *SS* = 1; Fig. 5C). For both marmosets, proportion correct and the standard deviation of their errors were approximately constant at these levels for signal strengths *SS* > 0.75 (corresponding to values of the stimulus range parameter < 90°). However, for both marmosets, proportion correct decreased (Fig. 5B) and the standard deviation of their errors increased (Fig. 5C) monotonically as signal strength was reduced.

In the absence of any coherent motion signal (i.e., signal strength, SS = 0), the error distributions of both marmosets were approximately uniform (Fig. 5A; bottom) and the proportion correct for both monkeys was consistent with chance performance (indicated by the dashed gray line in Fig. 5B). For this condition, the standard deviation of the Normal distribution of the mixture model exceeded 100° (Fig. 5C), again, consistent with chance performance.

The reduction in performance, both the decrease in proportion correct and the increase in standard deviation of the marmoset’s errors, as signal strength is reduced reflects the increasing difficulty of the task. This increase in difficulty is also reflected in the time taken for the marmosets to reach a decision (Fig. 6). Figure 6A shows distributions of reaction time for both marmosets over a range of motion signal strengths. These distributions are unimodal and highly skewed. To quantify changes in reaction time as signal strength was varied, we computed the median reaction time for all trials of each signal strength (Fig. 6B). For the strongest motion signal, the reaction times of both marmosets were similar (median, 253 and 259 ms, 95% CIs [250, 258] and [258, 260], for monkeys S and H, respectively). For both marmosets, reaction time increased monotonically as signal strength was reduced. In the absence of any coherent motion signal (*SS* = 0), median reaction times were 398 and 382 ms, 95% CIs [391, 408] and [374, 392], for monkeys S and H, respectively. Thus marmosets systematically trade speed for accuracy, integrating for longer time periods when noise or uncertainty in the sensory signal is higher.

### Comparison of motion estimation in marmoset and human observers

To compare motion estimation performance of the marmoset with that of human observers, we had four human subjects perform the same motion estimation task. As for the marmosets, we quantified human performance in terms of their proportion correct (Fig. 5B), and the precision of their choices (by fitting the mixture model; Fig. 5C). Over all signal strengths, the human observers performed better than the marmosets (Figs. 5B and 5C).

On trials containing strong motion signals, the human subjects performed significantly better than the marmosets: proportion correct 0.99, 1.0, 0.98 and 0.99, 95% CIs [0.98, 1.0], [1.0, 1.0], [0.96, 1.0] and [0.96, 1.0], for humans 1–4, respectively (Fig. 5B). Similarly, the human subjects’ choices were more precise, with standard deviations approximately half those of the marmosets (at the highest signal strength tested, *SS* = 0.875, the standard deviations of the human subjects’ perceptual errors were 10.6°, 9.9°, 11.9° and 13.4°, 95% CIs [10.0, 11.1], [9.5, 10.4], [11.2, 12.6] and [12.7, 14.1], for humans 1–4, respectively; Fig. 5C). Like the marmosets, the performance of all four human subjects decreased as the strength of the motion signal was reduced.

Human observers also showed a systematic increase in reaction time as signal strength was reduced (Fig. 6B), reflecting a similar trade-off in speed vs. accuracy to that seen in the marmosets. However, this trade-off was less dramatic in the human observers than in the marmosets (Fig. 6B). Notably, human observers were substantially slower than marmosets in the high signal strength conditions (median reaction times 296, 288, 290 and 321 ms, 95% CIs [296, 297], [287, 288], [288, 296] and [316, 323], for humans 1–4, respectively vs 250 and 258 ms, 95% CIs [249, 256] and [257, 259], for monkeys S and H, respectively; *SS* = 0.875, the highest signal strength seen by the human observers). However, this relationship was reversed under conditions of greater stimulus uncertainty (median reaction times 330, 365, 323 and 365 ms, 95% CIs [329, 331], [354, 379], [317, 325] and [362, 371], for humans 1–4, respectively vs 391 and 365 ms, 95% CIs [383, 400] and [357, 375], for monkeys S and H, respectively; *SS* = 0.125, the lowest signal strength seen by the human observers; Fig. 6B). Overall, marmoset and human observers exhibited similar qualitative trends for increasing reaction time with decreasing signal strength, but differed in the scale of this effect, with marmosets showing a greater dependence of reaction time on signal strength and less precision in their estimates.

### Choices reflect stimulus fluctuations

To assess whether differences in performance between humans and marmosets are due to a difference in their temporal integration strategies, we used psychophysical reverse correlation to estimate temporal kernels for both marmosets and the four human observers (see *Materials and methods*). We first aligned each trial with the time of motion stimulus onset and computed the correlation between the mean dot direction on each stimulus frame and the monkey’s perceptual choices, across all trials (Fig. 7A). The kernels estimated for the two marmosets were indistinguishable. Both marmosets exhibit small, but positive correlations over a period of approximately 250 ms following onset of the motion stimulus, with early frames having a modestly greater influence on their choices (Fig. 7B). Because there was no constraint in the task to wait for the entire stimulus duration prior to initiating the saccade, it is also useful to look at the temporal profile of integration prior to the saccade by realigning all trials to saccade onset (Fig. 7C). These kernels reveal that the marmosets integrate motion over a relatively brief time window that peaks approximately 150 ms before saccade onset and ends approximately 50 ms before the saccade. This latter period in which sensory integration contributes little to the perceptual decision is likely required to plan the eye movement. This delay is commonly observed in human behavior and is termed the saccade dead time (Findlay and Harris, 1984; Becker, 1991).

**Figure 7.**
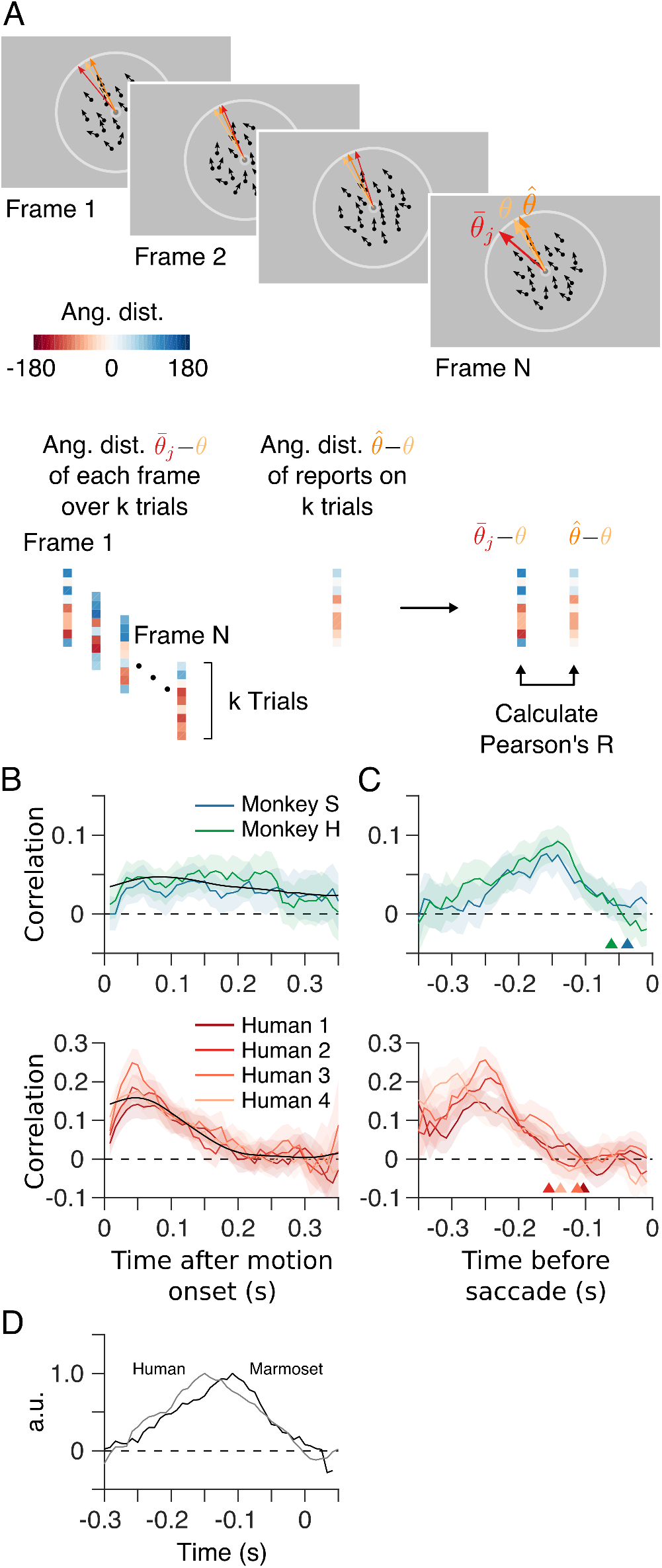
Choices reflect recent stimulus history. **A**. To assess the influence of different stimulus epochs (frames) on each subject’s choices we estimated their temporal integration weights (temporal kernel) using psychophysical reverse correlation. For each trial, k, we computed the difference between the mean motion direction over all dots, 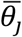, for each stimulus frame, j, and the target motion direction, *θ*_*k*_. For each frame, j = 1…N, we constructed a vector containing these differences for all trials. We then computed the correlation between this vector, for each frame, with the vector containing the difference between the subject’s choices, 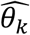, and the corresponding target motion directions, *θ*_*k*_, to reveal the subject’s ‘temporal kernel’. **B**. Temporal kernels for each subject for trials aligned with the onset of the motion stimulus. **C**. Temporal kernels for each subject as in B, after re-aligning each trial with the onset of the saccade indicating the subject’s choice. Arrow heads indicate the estimated saccade dead time for each subject. For comparison, B and C also show temporal kernels for four human observers performing the same motion estimation task. Shaded regions indicate bootstrap estimates of 95% confidence intervals. **D**. Average saccade-aligned temporal kernels for marmosets and humans, accounting for differences in saccade dead time and normalizing to the peak amplitude.

We considered to what extent the kernel locked to stimulus onset (Fig. 7B) could be explained by the kernel aligned to the saccade (Fig. 7C), after taking into account the variation in reaction times (Fig. 6). We first averaged the kernels for both marmosets shown in Figure 7C. For each trial the marmosets performed, we shifted this average kernel by the observed reaction time, in effect re-aligning this kernel with stimulus onset. We then averaged these shifted kernels across trials to obtain an estimate of the temporal kernel that would result from alignment with stimulus onset. This prediction is shown overlaid in Figure 7B (black line) and provides a good approximation of the kernels estimated by reverse correlation (Fig. 7B). Thus it appears that the contribution of motion information to marmoset decisions is most parsimoniously explained by a pre-saccadic integration process followed by post-decision saccade dead time.

We performed the same correlation analyses for the four human observers (lower panels in Figs. 7B and 7C). Temporal integration kernels for all four humans were remarkably similar to each other. In contrast to the marmosets, human kernels aligned to stimulus onset (Fig. 7B) exhibit a strong peak early in the stimulus presentation rather than the more prolonged and uniform kernels for the marmosets. However, estimates of the kernels aligned with saccade onset (Fig. 7C) are very similar to those for the marmosets, after allowing for a difference in saccade dead time. Human observers exhibit saccade dead times that extended out to 150– 200 ms prior to the saccade, as compared to the briefer 50–100 ms dead time for marmosets. As was the case for marmosets, it was possible to provide an accurate prediction of the stimulus aligned kernels for the human observers by shifting their average saccade-aligned kernel based on the observed reaction times (Fig. 7B, black line). The early peak observed for human subjects, but not marmosets, reflects the smaller variation in reaction times of the humans (median reaction time of the human observers varied by 47 ms, on average, over the range of signal strengths tested) compared to the marmosets (135 ms).

The temporal kernels estimated after aligning to the time of the saccade reveal that both marmosets and humans integrate over a short window of approximately 150–200 ms, followed by a period of little or no integration, presumably the dead time required to plan the eye movement. Saccade dead time (see *Materials and methods*) was considerably longer in humans (103, 155, 112 and 138 ms, 95% CIs [92, 135], [144, 170], [100, 125] and [126, 150], for humans 1–4 respectively) than in marmosets (37 and 61 ms, 95% CIs [0,67] and [38, 88], for monkeys S and H, respectively). To assess the similarity of the temporal kernels for marmoset and human observers, accounting for the differences in dead time, we re-aligned the kernels for each subject by subtracting their corresponding dead time, averaged these kernels across subjects and then normalized by the peak amplitude. The resulting normalized average kernels for marmosets and humans were highly correlated (Pearson’s R = 0.85, p < 0.001; Fig. 7D).

### Pre-saccadic eye drift reflects a less accurate read-out of stimulus motion

The random dot motion stimulus often evoked small drifts in eye position during its presentation (Fig. 8A). To determine if these movements were on average driven by the onset of stimulus motion we projected instantaneous eye velocity (see *Materials and methods*) onto the direction of the motion stimulus, and averaged the resulting eye speed signals across all trials for each motion signal strength (Fig. 8B). For both marmosets, these average eye speed traces revealed epochs of significant drift (speed > 0) beginning approximately 75 ms after onset of the motion stimulus. This latent period was longer, approximately 125 ms, in the human observers. Eye speed in the direction of the motion stimulus was greatest for coherent motion (*SS* = 1.0) and decreased systematically as motion signal strength was reduced (Fig. 8B).

**Figure 8.**
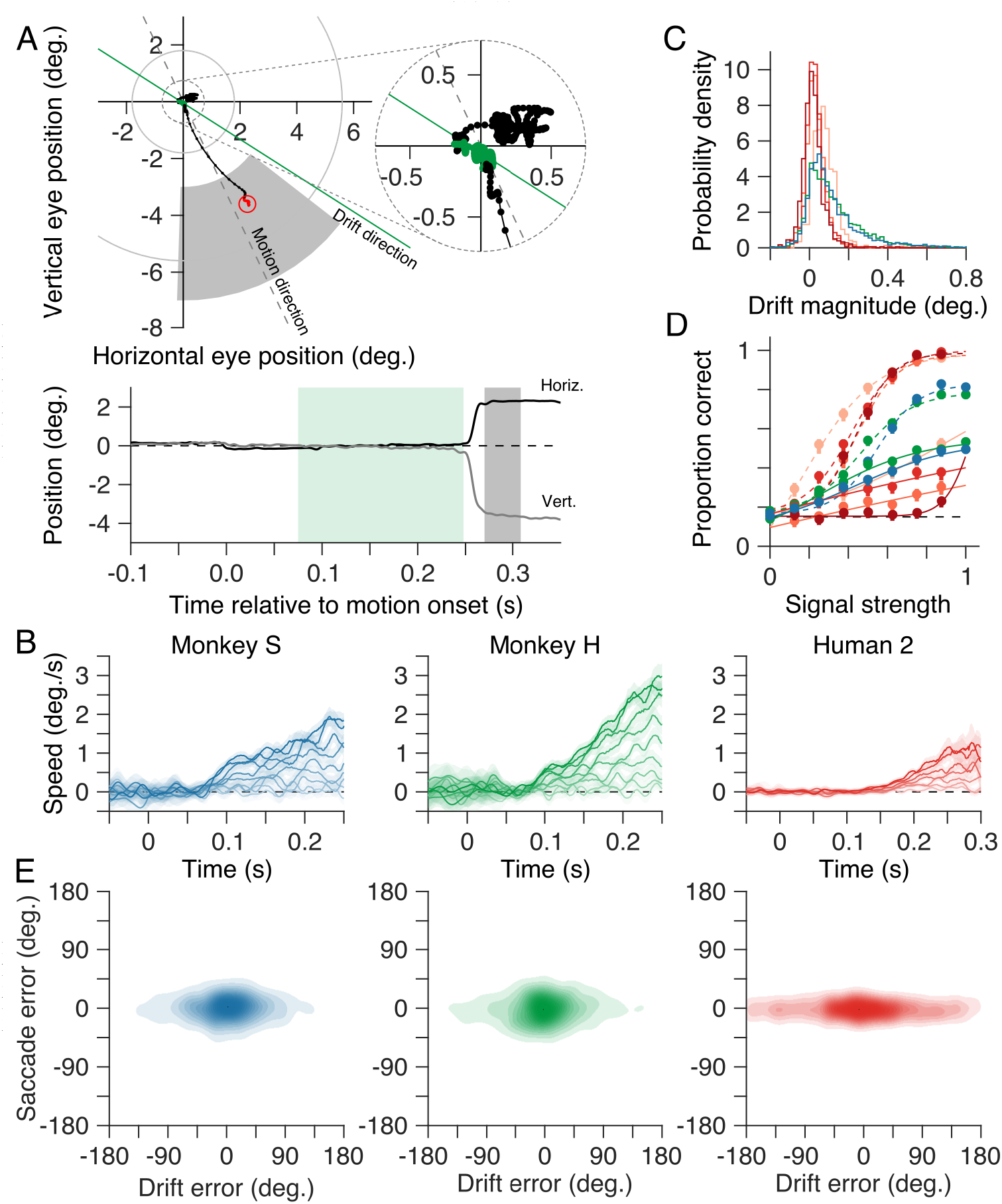
Choices are independent of small drifts in eye position during motion presentation. **A**. Horizontal and vertical eye position from a representative trial from one monkey (Monkey H, SS = 0) plotted in space (upper panel) and over time (relative to motion stimulus onset; lower panel). We often observed small drifts in eye position during presentation of the motion stimulus (green symbols in the upper panel and green shaded epoch in the lower position-vs-time panel). **B**. Eye speed vs time relative to motion stimulus onset, projected onto the motion stimulus direction and averaged over all trial for each signal strength. On average, the eyes remain stationary for ~75 ms after motion onset (longer, ~125 ms, in human observers) before drifting slowly until onset of the saccade indicating the subject’s choice. Drift speed decreased systematically as motion signal strength was reduced. **C**. Magnitude of the drift in the direction of the stimulus motion. **D**. Proportion of correct trials as a function of signal strength, based on the drift vector direction. Proportion correct for both marmosets and humans decreased as signal strength was reduced. To aid comparison, dashed lines show proportion correct for each subject based on their choices (reproduced from Fig. 5B). **E**. Density plots of saccade error vs drift error to assess the extent to which systematic drift in eye position could account for the subject’s subsequent choice. For both marmosets and humans, errors in the subject’s choices were independent of errors in drift direction. Same conventions as in Figs. 5–7.

We next quantified how well the pre-saccadic eye drift tracked stimulus motion on a trial by trial basis. We computed the magnitude and direction of drift on each trial by taking the vector sum of instantaneous eye velocity within a window beginning 75 ms (125 ms for the human observers) after motion stimulus onset and extending up until 20 ms before onset of the saccade which terminated each trial. These drift vectors exhibited small components in the direction of the motion stimulus (distributions shown in Fig. 8C; median 0.08 and 0.09 deg./s, 95% CIs [0.075, 0.083] and [0.085, 0.095], for monkeys S and H, respectively; *SS* = 1.0).

To assess the accuracy of drift as a measure of stimulus motion, we computed the drift error on each trial as the angular difference between the drift direction and the stimulus motion direction. To facilitate comparison with choices indicated by the saccades, we then computed the proportion of correct trials (error with the reward window; see *Materials and methods*) as a function of signal strength (Fig. 8D). For strong motion signals, the proportion correct for both marmosets based on drift direction (solid lines, Fig. 8D) was almost half that based on their saccades (dashed lines, Fig. 8D). At the highest signal strength, the proportion correct was 0.53 and 0.55, 95% CIs [0.49, 0.65] and [0.51, 0.63], for monkeys S and H, respectively. For the human observers, proportion correct based on the drift vector was worse than that of the marmosets (Fig. 8D). This is likely due to the humans’ better control of fixation compared to the marmosets. For both the marmosets and the human observers, proportion correct decreased as signal strength was reduced. In the absence of any coherent motion signal (i.e., signal strength, *SS* = 0) proportion correct was consistent with chance performance.

Last, we considered to what extent the pre-saccadic drift might influence the subsequent saccade choices. If the drift influenced subsequent saccade choices, then we would predict that trial by trial errors of the two measures would be correlated. However, if drift and saccade choices were independently driven by the motion stimulus, we would expect the errors to be uncorrelated. We computed the correlation between errors in the monkey’s choices (revealed by their saccades) with the corresponding errors in the drift vectors (Fig. 8E). We found no significant correlation, for either monkey, even at the highest signal strengths tested (R = −0.01 and 0.01, 95% CIs [−0.05, 0.03] and [−0.04, 0.07], for monkeys S and H, respectively; Fig. 8E). Similarly, we found no significant correlation for the human observers: R = 0.0, −0.01, 0.08, −0.08, 95% CIs [−0.07, 0.08], [−0.08, 0.05], [−0.01, 0.15] and [−0.24, 0.07], for humans 1–4, respectively. This suggests that while the drift is driven by the motion stimulus, it does not directly influence the saccade choice.

## Discussion

Marmoset monkeys promise to offer unprecedented access to circuit-level measurements in a primate brain during visually-guided behaviors (for review, see Mitchell and Leopold (2015)). Here, we developed a paradigm for studying motion perception in the marmoset that is amenable to their oculomotor behavior and to the study of neural population codes. We found that marmosets learned the motion estimation task within a reasonable timespan and performed with an accuracy that, although short of human performance, was qualitatively similar. Both marmosets and humans had accuracy and reaction times that depended on the difficulty of the task. Additionally, both species’ reports depended systematically on temporal fluctuations in the direction of the stimulus, indicating brief temporal integration. Marmosets and humans used similar temporal weighting of evidence that favored information immediately preceding saccade onset, differing primarily in the duration of their saccade dead time. Both marmosets and humans also exhibited low gain eye drift, during fixation, that follows stimulus motion with short latencies from stimulus onset. This low gain drift was less accurate as an estimate of stimulus motion direction than the subsequent saccade choices, and was not predictive of errors in the subsequent saccades. This suggests that while eye drift and saccade choice are both related to stimulus motion, for both marmosets and humans they may rely on partly non-overlapping mechanisms.

Estimation paradigms, like the one employed here, offer several advantages over traditional two-alternative forced choice tasks. The primary advantage is that they produce a distribution over perceptual reports, which can more accurately distinguish the computations that are involved in generating the percept (Webb et al., 2007; Webb et al., 2011). This extra level of detail can distinguish between perceptual algorithms (Webb et al., 2007; Webb et al., 2011; Laquitaine and Gardner, 2018) or may reveal dynamic biases (Panichello et al., 2018) in a manner that is not possible with forced choice paradigms. Additionally, estimation tasks can target all stimulus features (e.g., direction) equally, making them better suited for large population recordings from neurons with diverse tuning, as opposed to discrimination tasks, which are typically optimized for a few neurons under study. The estimation task employed here did not require long training periods for the marmosets (approximately 3 months). For comparison, on a two-alternative forced choice motion discrimination task in macaque monkeys, Law and Gold (2009) reported that while performance plateaued quickly for one monkey (< 20 sessions), the second monkey required considerably more training (>60 sessions). Our marmosets learned the motion estimation task (for high signal strength stimuli) in 40–60 sessions. One caveat with estimation paradigms is that they produce fewer repeats of each condition. For comparisons to physiology, it is possible to reduce the number of stimulus difficulties presented per session, or pool across sessions using chronic recordings. However, we believe that model-based approaches to comparing behavior and physiology will be sufficiently rich, specifically because motion-perception is such a well-developed field.

One distinctive feature of our estimation paradigm is that observers were free to indicate their choice while the stimulus presentation was ongoing. This may have important consequences for how we interpret the accumulation of motion evidence. We found that motion integration was better explained in a saccade aligned reference frame rather than aligned to stimulus onset. Both human and marmoset observers integrated motion over a short window of approximately 150-200 ms, followed by a period of little or no integration, presumably the dead time required to plan the eye movement. We could reliably predict the motion integration that was locked to stimulus onset based on the kernels aligned to saccade onset, but the reverse was not true. If the observers had been constrained to maintain fixation until completion of the stimulus epoch, we would not have been able to examine these dynamics, and may have erroneously concluded that humans preferentially integrate earlier evidence in the stimulus epoch whereas marmosets do not. A recent study in which macaque monkeys performed a motion discrimination task also allowed subjects to indicate perceptual choices prior to the end of the stimulus epoch (Okazawa et al., 2018). They found comparable trends in the weighting of motion evidence, with peaked response-aligned kernels followed by a delay – the saccade dead time – preceding the saccade (Okazawa et al., 2018). Similar to the kernels we observe, Okazawa et al. (2018) found that kernels estimated when trials were aligned to stimulus onset exhibit an apparent early weighting of sensory evidence. They showed that this pattern is nonetheless consistent with a decision making process that involves an integration process that weights sensory evidence equally over time, provided that there is an accumulation to bound followed by a variable dead time preceding the saccade response. There are, however, other possible models that could explain the weighting of sensory evidence which might also apply to our data (Yates et al., 2017; Levi et al., 2018). Our results remain agnostic to the underlying mechanism, but do support the notion that both marmosets and humans use a similar temporal weighting of evidence, differing primarily in the duration of their saccade dead time.

Although both marmosets and humans showed a dependence of speed and accuracy on stimulus strength, reaction times varied over a relatively short range compared to prior work on perceptual decisions (Palmer et al., 2005). Our reverse correlation analysis confirmed that both humans and marmosets exhibited relatively short integration windows on the order of 150–200 ms. This appears shorter than the integration windows observed for alternative forced-choice decision-making paradigms (Hanks et al., 2011; Kiani et al., 2014; Okazawa et al., 2018), more closely resembling reaction times for perceptual and pursuit tasks (Osborne et al., 2007; Price and Born, 2010). The relatively short integration windows for our estimation paradigm make it amenable to neurophysiological study using short counting windows. For example, when computing choice correlations and other measures of decision making, there would be value in being able to localize the relevant motion integration to a relatively short epoch. Moreover, the short integration windows we observe seem likely to be more relevant to the natural timescales of behavior in which perceptual decisions are normally made.

Marmosets also exhibited small drift movements during fixation along the direction of stimulus motion that occurred just before their saccade choice. Although our paradigm discouraged eye drift due to the fixation constraint, these movements persisted over training with very low velocities, typically under 20% gain for marmosets and 10% gain for humans. We were able to average the velocity traces across trials to resolve their time course relative to motion onset. Average eye velocity began to ramp up along the stimulus motion direction at latencies of 70-80 ms in marmosets and 100-150 ms in humans. This low gain following of stimulus motion resembles involuntary ocular following movements in tasks using wide-field motion stimuli with no fixation constraint (Miles et al., 1986; Gellman et al., 1990). We quantified the accuracy of the eye drift direction from motion onset up until the saccade choice as a report for the motion direction but found it was poor relative to the saccade choices made by the subjects (Fig. 8D). The errors in eye drift direction did not correlate well, on a trial by trial basis, with errors in the saccade choices (Fig. 8E), suggesting that although both are driven by motion information in the stimulus, they may rely on at least partly different neural mechanisms. A previous study has also reported some degree of independence for motion read-out in perception as opposed to ocular following in human participants (Simoncini et al., 2012; Glasser and Tadin, 2014; Price and Blum, 2014). It is possible that eye drift might provide a more accurate measure of motion direction under conditions different to those in our study. We optimized our task to focus on saccade choices, and by using a fixation point, discouraged smooth following of stimulus motion. Thus, it seems likely that more accurate following responses would occur using conventional ocular following paradigms.

The current paradigm employed random motion stimuli presented at foveal and parafoveal eccentricities rather than in peripheral apertures. Classical studies in macaques have normally used peripheral stimuli that were localized precisely to match the receptive fields of the neurons under study in daily sessions (Newsome et al., 1989; Britten et al., 1992). This carries a clear advantage for correlating the activity of those neurons with behavioral choices, as they will be the most relevant to the behavior. It should be possible to adapt the current paradigm for peripheral stimulus locations (Nichols and Newsome, 2002), but the use of foveal stimuli may also carry complementary advantages for large-scale array recordings in marmosets. First, when using arrays, it is not necessarily optimal to tailor stimuli for individual neurons. In that context, targeting foveal locations may be desirable. Besides the prominence of the fovea in primate vision, several recent studies demonstrate selective weighting of foveal information for motion integration in smooth eye movements (Mukherjee et al., 2017). Further, it is relatively straightforward to reliably target foveal representations in areas MT and MST of the marmoset because they are readily accessible at the cortical surface. In short, while the use of stimuli near the fovea in the current behavioral design deviates from several previous studies, it may confer distinct advantages for studying neural coding in the marmoset.

Recent studies in anesthetized marmosets have been able to span foveal and peripheral representations within MT/MST using planar silicon arrays (Chaplin et al., 2017; Zavitz et al., 2017). Similar methods could be readily employed in awake marmosets in conjunction with the current behavioral paradigm, and would enable us to examine the neural code for motion perception from large-scale populations. Thus, our findings establish motion perception in the marmoset as a highly promising model system for studying the population codes and circuit mechanisms that underlie perception in a primate species.

